# Variation in survival and growth following prolonged darkness in a polar diatom species

**DOI:** 10.64898/2026.02.02.703299

**Authors:** Patricia Mrazek, Sinéad Collins

**Affiliations:** University of Edinburgh

**Keywords:** Keywords diatom, polar night, genotype sorting, natural selection, population growth, Southern Ocean

## Abstract

- Phytoplankton are the major primary producers in the Southern Ocean, participate in the global carbon cycle, nutrient cycles, and are at the base of the food-web. These polar ecosystems are unique in their extended periods of darkness in the winter.
- Prolonged darkness has the potential to exert selection that affects the composition of diatom communities if there is differential survival of diatoms in the dark, variation in population growth rates in subsequent light periods, or both.
- We tested whether prolonged darkness has the potential to exert within-species selection on a model polar diatom species by exposing 5 strains of the polar diatom *Porosira glacialis* to prolonged darkness at two different temperatures in the laboratory. We measured population survival in the dark, growth rate upon re-illumination, and between strain variability in these traits.
- We found a pronounced decline in survival and growth rate with time spent in the dark, as well as important intraspecific variation in these.
- Higher temperature exacerbated declines in growth and survival.
- Our results show that the darkness of polar night can exert selection within diatom species, with implications for phytoplankton community composition and subsequent impacts on Southern Ocean biogeochemical cycles.

## Introduction

The Southern Ocean (SO) is an extremely productive habitat, fuelled by photosynthetic microbes (phytoplankton). Diatoms are the most abundant group of phytoplankton in the Southern Ocean and make up over 40% of phytoplankton biomass there (Kale and Karthick, 2015), where they also play an important role in carbon uptake, biogeochemical cycles, and polar foodwebs. The SO is disproportionately important in global biogeochemical cycles, given its outsized contribution to biological carbon uptake and underlying nutrient cycling processes (Armbrust, 2009). Nutrient export (N, P, Si) from the Southern Ocean accounts for over to 60% of nutrients exported to temperate and tropical latitudes (Sarmiento *et al*., 2004; Fripiat *et al*., 2021), and impacts global primary production, ecosystems and fisheries. Since these processes are affected by the composition of diatom communities, understanding how selection shapes both inter- and intraspecific variation in polar diatoms is fundamental to understanding polar ecosystems.

A key driver of primary productivity is light (Arteaga *et al*., 2020), and springtime productivity in polar systems is dependent on the end of the polar night, where phytoplankton populations have just experienced a prolonged period of darkness that can reach up to 4 months depending also on ice and snow cover (Biggs *et al*., 2019; Young *et al*., 2024). Thus, polar diatoms must both survive prolonged periods of darkness, and resume population growth upon re-illumination. Variation in dark survival and subsequent population growth in the light has the potential to drive both differences in strain composition within species (and thus species average trait values) and the species composition of wider diatom communities (Fang and Sommer, 2017; McMinn *et al*., 1999; Reeves *et al*., 2011) (Figure 1). These changes in species and community composition have potential effects on polar food webs and biogeochemical cycling if functional traits, such as cell size, covary with overwinter survival or spring growth (Biggs *et al*., 2019). These shifts in diatom community composition can then affect rates of nutrient uptake and export, as well as the food value of diatoms for grazers (Deppeler and Davidson, 2017; McMinn *et al*., 1999, Litchman, 2000).

**Figure 1.**
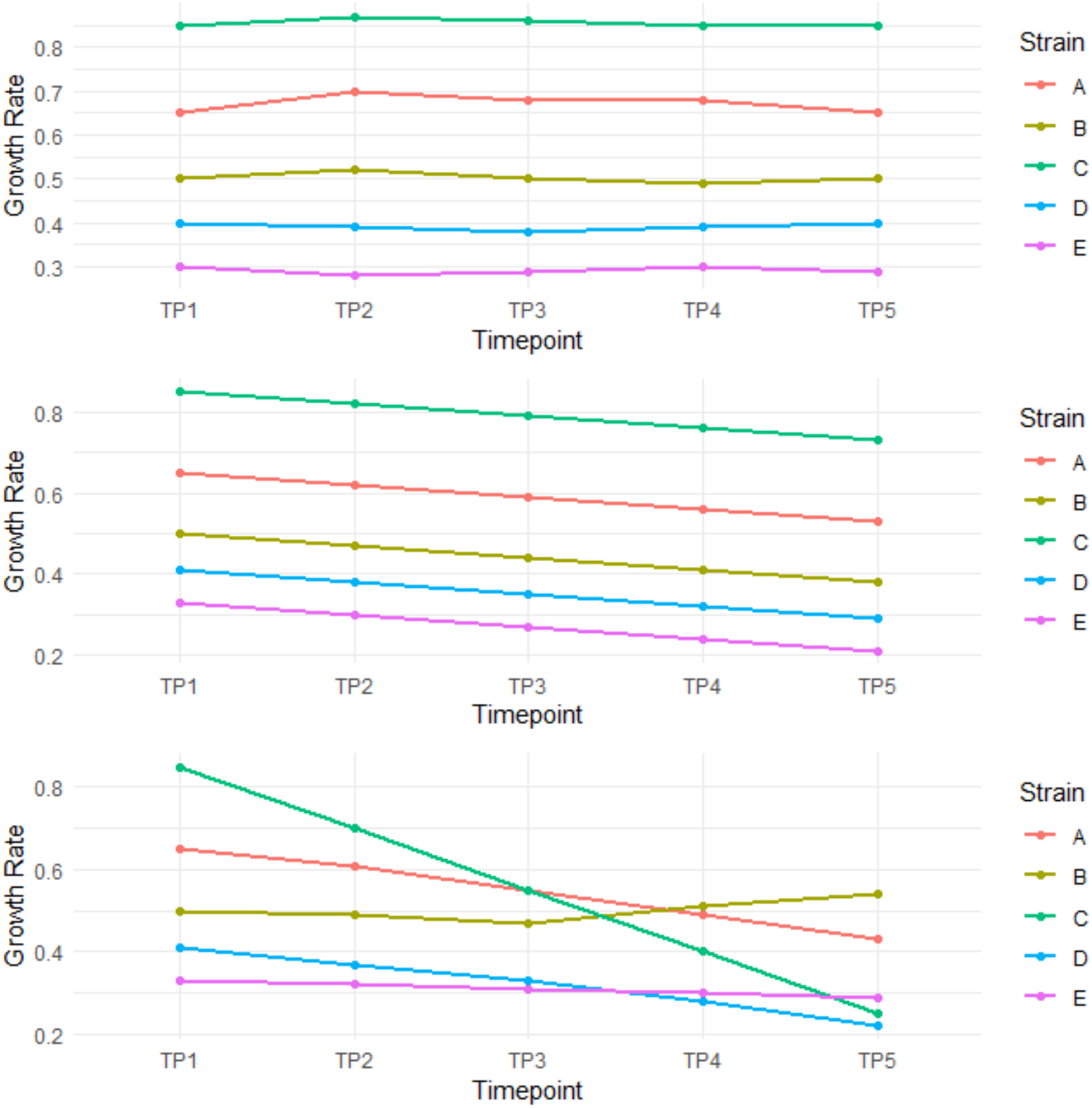
Conceptual figure of the possible growth rate responses of several strains to overwintering. Growth rate might remain approximately stable (a), it might decrease at equal rates in all strains with time spent in the dark (b), or different strains’ growth rates might respond differently to overwintering, resulting in a rank-order change (c).

Information on how diatoms survive weeks or months of very low or no light is rapidly growing. Diatoms and other polar phytoplankton have specific traits and strategies linked to surviving polar nights. These strategies include reducing metabolism, sexual reproduction and spore formation, and maintaining high photosynthetic efficiency to resume photosynthesis immediately once light becomes available (Armbrust *et al*., 2009; Sciandra *et al*., 2022). Recent evidence from the polar diatom *Fragilariopsis cylindrus* suggests that diatoms reduce their metabolic rates and rely on lipid stores to survive dark periods (Joli *et al*., 2024). In addition to surviving periods of darkness a key strategy in diatoms’ success at polar latitudes is their ability to resume photosynthesis almost immediately upon re-illumination, even at very low light intensities (Morin *et al*., 2020; Reeves *et al*., 2011). Maintaining the ability to photosynthesize during extended darkness gives polar diatoms a competitive advantage once light becomes available (Hoppe *et al*., 2024; McMinn *et al*., 1999). These and other mechanisms, reviewed in Berge *et al*. (2015), are painting an increasingly detailed picture of how phytoplankton survive through the polar night, and photosynthesize immediately following it. However, there is a lack of data on variation in dark survival and subsequent photosynthesis-driven population growth, which has the potential to affect the composition of diatom communities (Fang and Sommer, 2017).

Studies to date have focused on how single strains survive in the dark. A detailed study by Joli *et al*. (2024) shows how a wide range of physiological traits change during periods of darkness in a single strain of *Fragilariopsis cylindrus*, but does not address growth following the dark period. van de Poll *et al*. (2020) explore the relative survival of two Polar diatom species (*Thalassiosira antarctica, Thalassiosira nordenskioeldii*) and two flagellate species (*Rhodomonas sp*., *Micromonas sp*.) over an 8-week darkness period. They find that the diatoms have a higher survival than flagellates, which is consistent with diatoms having a competitive advantage once light becomes available. They also report differences between the two diatom species *T. antarctica* and *T. nordenskioeldii*, but lack the statistical power to explore this variation in more detail.

In addition to variation in dark survival and subsequent population growth within and among species, there is variation in the length of dark periods that populations experience. This is due to differences in latitude and depth, as well as the timing and extent of ice formation and melt (Gordon, 1981). The latter of which is rapidly shifting in polar regions (Gordon, 1981; Lasternas and Agustí, 2010; Young *et al*., 2024), as warming water and atmospheric temperatures are driving earlier ice-melt and later ice-formation, with lower overall ice coverage and thickness (de la Mare, 2009).

In this study, we examine how biological and environmental variation affect diatom dark survival and subsequent growth in the light, using five strains of the common Antarctic diatom *Porosira glacialis*. Single-strain populations were placed in the dark for up to three months at two different temperatures (well below, and near, T_opt_ for these strains). Over three months, they were subsampled five times, and at each sampling time we measured cell survival and population growth rates upon reillumination. We found that the proportion of viable cells in each population declines with the length of time spent in the dark. When populations are re-exposed to light, population growth rate also declines with the length of time spent in the dark. These trends were amplified at the elevated temperature. We find substantial intraspecific variation in survival and growth responses at both temperatures, suggesting that variation in dark survival and growth following dark survival can contribute to strain sorting within polar diatom species.

## Materials and Methods

### Strain Collection and Cultivation

The diatoms used for this project were isolated from the Southern Ocean during a single cruise in 2017. Each strain was cultured from a single isolated cell, which ensured genetic homogeneity of each strain (Bishop *et al*., 2022). The collected strains were continuously cultured since collection at 2C under 24h light, with regular transfers into fresh F/2 media (Guillard 1975). In preparation for the experimental treatment, 5 strains (Pg590, Pg595, Pg633, Pg654, Pg716) of the diatom *Porosira glacialis* were acclimated to temperatures of 1°C (low) and 4°C (high) respectively, and +-30 micromolar light intensity, for 5 days prior to the beginning of experimental treatments. All cultures were actively growing at the beginning of the experiment.

### Experimental Design

This study was designed to measure variation in survival and growth attributable to intraspecific variation and length of dark period. This was done at two temperatures, below and near the optimal temperature for growth for this species (Bishop *et al*., 2022). To accommodate the replication needed, culture size was limited, which allowed the growth and monitoring of 60 total cultures. All populations were grown as single strain cultures. Experiments were initiated from cultures of approximately equal cell density for dark treatments (3500cells/ml, +-1000 cells/ml) and light treatments (350cells/ml, +-100cells/ml) (see S.I. Table S1), with 3 replicates per strain per treatment. Populations were grown in non-vented culture flasks in 50mL of F/2 media (Guillard 1975). Dark treatment flasks were wrapped in 3 layers of tin foil. Three replicates of each strain-by-treatment combination were maintained in respective high or low temp incubators for the duration of the experiment. For a diagram of the experimental setup see figure S1.

Each culture was sampled approximately every 14 days for mortality assays for 2 months, then at the end of 3 months of treatment. Samples from each culture were re-inoculated into light conditions approximately every 10 days for the duration of 2 months, then after 2 months (see exact timeline in S.I. Appendix 2, Table 2). The population growth rate of the re-illuminated samples were measured to assess culture viability. At the end of the 3 month experiment, we also measured cellular polar lipid content using flow-cytometry. The precise experimental design, set-up and timeline can be found in the S.I. Appendix 2, Figures S1-S3.

### Mortality Assays

Mortality was assessed by counting live and dead cells from a sample of each culture and calculating the proportion of dead cells present. Cells were stained with 2% Evans Blue stain, as in Samuels *et al*. (2019), adapted for this species by using a staining time of 1 hour, +-15 minutes. Samples were concentrated 15-fold by centrifugation in a Eppendorf Centrifuge 5424R (8 minutes, 800rcf, 2C), mounted onto glass slides and imaged under an Olympus BX43 inverted microscope. Live cells showed up as pale green or green-gold, while dead cells were stained light grey-blue to dark-blue.

### Viability Assays

Viability post dark-treatment was assessed through population growth rates. An aliquot of each culture was inoculated into a well on a 24-well plate with fresh F/2 media, to a total volume of 1.9ml. Initial cell densities were measured by fixing cells with Lugol’s iodine (Utermöhl, 1958), concentrating 15-fold through centrifugation and counting cells using Sedgewick Rafter or Neubauer chambers (Serfling, 1949).

Post-dark cultures were grown at the temperature treatments they originated from, at 30 (+/-5) micromolar light intensities. Once cell growth was confirmed in all wells through chlorophyll fluorescence readings on the TECAN Spark BS050711 cell densities after growth were measured using counts as above.

From cell concentrations, the growth rate *μ* was calculated as

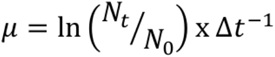

Where Nt is the cell concentration at time of harvesting, N0 is the cell concentration at time of inoculation and Δt-1 is the difference in days between the day of inoculation and day of harvesting. Finally, growth rates were corrected by the expected proportion of live cells at each timepoint, as calculated from the Evan’s Blue live/dead proportions.

### Lipid Content Assays

Following 3 months of experimental treatment, 5ml aliquots of each culture were sampled and stained with Bodipy dye, which reacts with polar lipid stores of phytoplankton cells (Park *et al*., 2025). The fluorescence of each sample was measured using flow cytometry. Chlorophyll fluorescence and relative sizes were measured by flow cytometry at the same time. At least 1000 cells of each sample were measured to estimate median per cell lipid and chlorophyll fluorescence.

### Statistical Analysis

Mortality data was analysed through logistic regressions, using the following model:

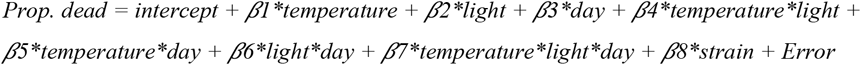

Growth rate was analysed through a linear regression with the following structure:

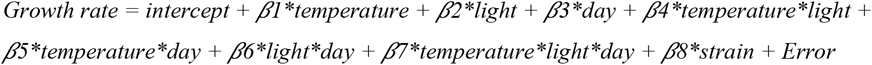

R and R Studio were used for all statistical analyses (R Core Team, 2024), data formatting and manipulation utilised the packages dplyr (Wickham *et al*., 2023) and tidyr (Wickham *et al*., 2024). Statistical analyses used packages lme4 (Bates *et al*., 2015) and lmerTest (Kuznetsova *et al*., 2017). Graphs were created using ggplot2 (Whickham, 2016)

### Lipid Analysis

Polar lipid content was measured using flow cytometry to detect Bodipy fluorescence (Park *et al*., 2025). From the gated population we extracted the Median and SD fluorescence values for Bodipy fluorescence using FlowJo software version 10.10.0 (BD Life Sciences). Gating information is in S.I., Appendix 3.

The role of lipids in cell mortality and viability was analysed using mixed regression models to address four specific hypotheses:

H0 - lipid content does not explain variation in mortality/growth rate upon re-illumination. H1 - initial lipid content best explains variation in mortality/growth rate. H2 - Final lipid content best explains variation in mortality/growth rate. H3 - Lipid loss best explains variation in mortality/growth rate.

## Results

### Phytoplankton Mortality

Because differential survival has the potential to impose selection during periods of darkness, we directly measured variation in mortality over time in our study. Overall, cell mortality was higher in the dark and increased with time; this effect was intensified at the warmer temperature. All strains followed this qualitative trend. Over all strains, we found a significant increase in dead cells in the dark over time at both temperatures (Z = 4.08, p < 0.001). Over all strains, cells in the dark were approximately twice as likely to be dead than in the light at any given timepoint and treatment (Odds ratio = 1.97, Z = 6.56, p < 0.001). This is consistent with wider research that indicates elevated mortality in the dark (Handy *et al*., 2024; Joli *et al*., 2024; Morin *et al*., 2020; Murphy *et al*., 2012; Reeves *et al*., 2011). We found a significant increase in dead cells in the dark over time (Z = 4.08, p < 0.001). Timepoint 5 showed increased mortality in the light treatment that is likely an artifact, as mortality is usually steady in standard cultures maintained under standard conditions. Analysing our data with and without timepoint 5 did not change the trends we report.

There was important variation in mortality between strains. When fitted as fixed effects in our model, each strain had a p value < 0.05. Strains Pg590 and Pg654 had consistently lower mortality, while Strains Pg595 and Pg716 had highest overall mortality. Strain Pg633 has the highest increase in mortality over time in the dark, in particular at 4°C.

### Growth rate upon re-illumination

We measured population growth rate upon reillumination after different times spent in the dark. Overall, population growth decreased after being maintained in the dark relative to cultures maintained in the light at the same temperature. (t = −5.158, p < 0.001, DF = 288). Populations maintained at the higher temperature (4°C) grew faster than cells maintained at 1°C (t = 5.714, p<0.001, DF = 288). However, growth rate decreased more rapidly over time in populations maintained at 4°C than at 1°C (t = −3.009, p = 0.003, DF = 288) in both dark and light treatments. The decrease in growth rate in the light treatment between timepoints 4 and 5 is likely an artifact as above and excluding this datapoint does not alter the overall trends of our data.

Population growth rate upon re-illumination varied between strains. This effect was driven by a single strain, Pg 716 (t = −4.989, p < 0.001), and over both light treatments, strain explained little of the variance in growth rate (∼5% overall, and ∼7% in the dark treatment), with most of the variance in growth upon re-illumination being due to the light treatments themselves (∼23%).

Considering both our results for cell mortality and growth rate upon re-illumination, we found that both survival and growth rate upon re-illumination decreased significantly over time spent in the dark. Strain variation attributable to differences between strains was more important in mortality responses than in growth rate responses.

### Lipid content

Polar diatoms are unusually lipid-rich, which is put forward as a partial explanation of their ability to survive long periods of dark or low light (Joli *et al*., 2024: Juchem *et al*., 2023; Zhang *et al*., 1998). To test whether variation in polar lipid content explained survival at this taxonomic level, we measured polar lipid content in our populations at the end of 3 months in the dark at both temperatures. Though being maintained in the dark depletes intracellular polar lipid stores (F = 440.3932, p << 0.001, DF = 56) (see S.I. Appendix 4), as expected, lipid stores correlated poorly with both mortality and growth rate in this experiment for populations maintained in the dark. However, variation in lipid content in the dark in this experiment was very low, suggesting that variation between, rather than within, species is more likely to underlie lipid-related mortality effects in polar diatoms.

For all results below, only populations maintained in the dark, and mortality and population growth upon reillumination at the final timepoint are considered, so that the below discusses how the absolute amount or concentration of polar lipids explains variation in dark survival and growth only.

We considered the proportion of dead cells as a response of median lipid concentration, strain and temperature, including an interaction between lipid concentration and strain. This model best captured the variation in the data, with an R2 of 54%.

Even when including median lipid concentration per cell in our analysis, temperature was the most important predictor of mortality (t = 4.470, p < 0.01), with an increase in mortality of 5.9% per degree and of 17.7% between the 1°C and 4°C treatments on average. We also found that strains differed in their baseline mortality within particular strains when median lipid concentration was included as a predictor (Pg595: t = −2.350, p = 0.029724, Pg654: t = −1.751, p = 0.096154, and Pg716: t = −2.004, p = 0.059531). Finally, our model also suggests some variability in the mortality response to decreases in lipid concentration depending on strain.

Interestingly, we found that lipid concentration, calculated as average lipid content per median cell spherical volume, may affect mortality at the final time point. Given the relatively low power of this study, we cannot discount that lipid concentration may have an effect on cell mortality, with our results suggesting that higher lipid concentration leads to a slightly lower proportion of dead cells (mortality) (effect size = −32.325, t = −1.986, p = 0.061682) Given the statistical effect at even a very small study size, this association requires more study.

We also examined whether lipid content explained variation in growth rate upon re-illumination. Variation in absolute lipid content or lipid concentration in cells maintained in the dark did not explain variation in growth rates upon re-illumination at the end of the experiment. A model fitting a three-way interaction of cell size, strain and temperature best explained the variation in the data (adjusted R2 = 79%) (see S.I. Appendix 3). Here, size is an important predictor of growth rate upon reillumination (t = −3.454, p = 0.00618), with smaller cells having faster growth rates (Estimate = 0.000003597 increase in growth rate per decrease in unit size). Temperature is another important predictor of growth rate upon re-illumination, with a 1°C increase in temperature associated with a 0.2532 decrease in growth rate upon re-illumination (t = −2.789, p = 0.01916, DF = 1).

This statistical model detects variation between strains. Here, strain Pg716 has a baseline growth significantly lower than other strains (using strain Pg590 as the reference strain), with baseline growth 0.435µ lower than strain Pg590 (t = −2.622, p = 0.02552, DF = 4). This model further indicates that strains Pg654 and Pg716 respond differently to an increase in temperature than other strains. The overall trend is for growth rate decrease with time spent at the higher temperature in the dark, these two strains have less of a decrease in growth rate over time spent in the dark at the elevated temperature (strain Pg654: t = 3.643, p = 0.00453; strain Pg716: t = 2.613, p =0.02589).

## Discussion

In this study we investigated intraspecific variation in mortality and growth rate upon re-illumination of 5 strains of *Porosira glacialis* subjected to different durations of darkness at two temperatures. We found that mortality increased with time spent in the dark, while population growth rate upon re-illumination decreased. We found significant intraspecific variation in both mortality in the dark and growth rate upon re-illumination, suggesting that both dark survival and growth after dark have the potential to shape diatom populations through the action of natural selection.

Our key result is that there was variation amongst *P. glacialis* strains in their mortality. We found that mortality increased over time in the dark in our experiment, where *P. glacialis* cells in the dark are almost twice as likely to be dead than in the light and the odds of a dead cell increase over time spent in the dark. Each strain displayed a different mortality. At 1°C strain Pg595 consistently showed the highest mortality, while strains Pg654 and Pg590 had just over half as much mortality by the end of the 3-month dark period as Pg595. Joli *et al*.’s (2024) recent study elucidated the physiological processes by which the polar diatom *Fragilariopsis cylindrus* can survive polar nights, by entering a state of hypometabolism, with respiration, consumption of cellular lipid stores and autophagy providing the small amounts of energy required by the cells in this state. The higher mortality, and variation in mortality in our study is consistent with variation in the exhaustion of cellular stores, degree of autophagy, an inability to achieve a state of quiescence, or a combination of these. Previous studies have found minimal mortality during periods of darkness in polar diatoms in laboratory experiments in different species, under differing culture conditions than used in our study (Morin *et al*., 2020; Reeves *et al*., 2011). The differences between our results and others points towards the need to better understand first, intraspecific variation in responses and second, how specific culture conditions affect the baseline mortality of cultures.

All populations were viable, in that they were able to grow, when reintroduced into the light after a period of darkness, but their population growth rates decreased with time spent in the dark. Viability has generally been assessed using photosynthetic efficiency, and many of these studies report high photosynthetic efficiency following prolonged darkness (Joli *et al*., 2024; Morin *et al*. 2020; Reeves *et al*., 2011). The logic of linking photosynthetic efficiency with viability is the presumed ability to resume growth if the photosynthetic machinery is working, and does not make predictions about how fast a population should resume growth after reintroduction into the light. In our study we focused on population growth rate upon re-illumination to understand how a key fitness component (population growth rate upon reillumination) varies with time spent in the dark. As in previous studies (Reeves *et al*., 2011; Martin *et al*., 2012; van de Poll *et al*., 2020; Hoppe, 2022), we found that all strains were able to resume growth following 90 days of darkness. However, the growth rate upon re-illumination decreased over time in the dark by an average of 0.0001329*μ* per day (Fig 3). Strain identity explained a small amount of the total variance in the growth rate model (5%-7%). Most of this strain effect is attributed to strain Pg716 which might indicate that certain strains can have differing demographic responses to darkness even within the same species. Even though this is a relatively small effect in our model, it is large enough to affect the strain composition of a species. For example, if the same number of cells of strain Pg590 and strain Pg716 were viable upon reintroduction into the light following 60 days of darkness, strain Pg590 would have a population size 1.55 times larger than strain Pg716 after 7 days under our growth conditions. Differences in how strains of the same species of polar diatom survive prolonged periods of darkness (Fig 2) suggests that the polar night can act as a strong selective event that shapes polar diatom communities (Juchem *et al*., 2023; Litchman, 2000; Wietz *et al*., 2021). The importance of the polar night as a selective event is further increased by strain-specific growth rates following various durations of darkness (Fig 3). Therefore, combining the effects on survival and viability indicates the potential for large differences in diatom community composition between autumn and spring in the Southern Ocean.

**Figure 2.**
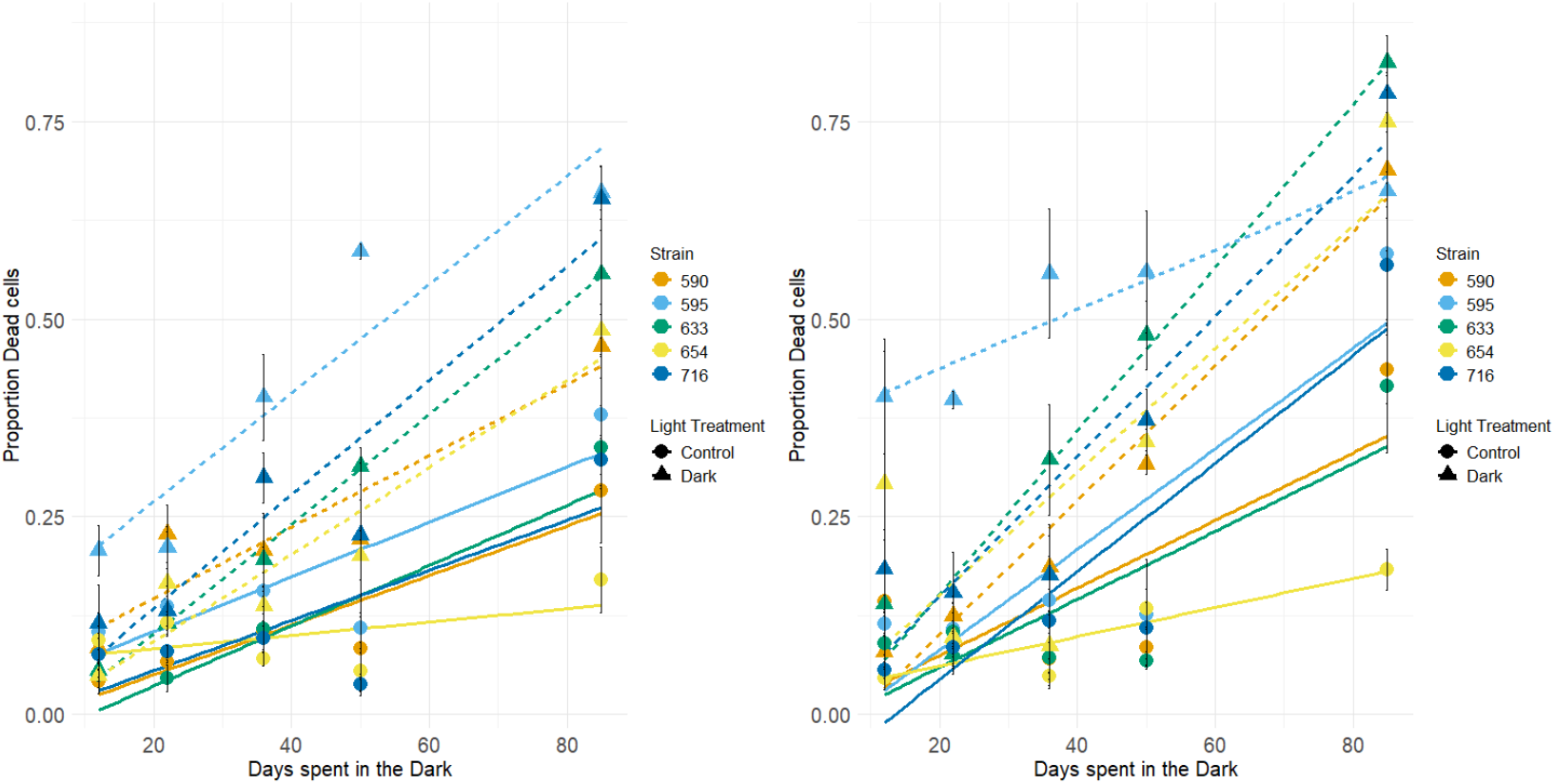
Proportion of dead cells over time of both light and temperature treatments. a) shows the proportion of dead cells in control-light cultures (round points and solid lines) and dark cultures (triangles and dashed lines) in the 1°C treatment. b) shows the proportion of dead cells in control-light cultures and dark cultures in the 4°C treatment. Black bars at each point correspond to standard error at each measurement. The colours of each line correspond to the 5 different strains of Porosira glacialis.

**Figure 3.**
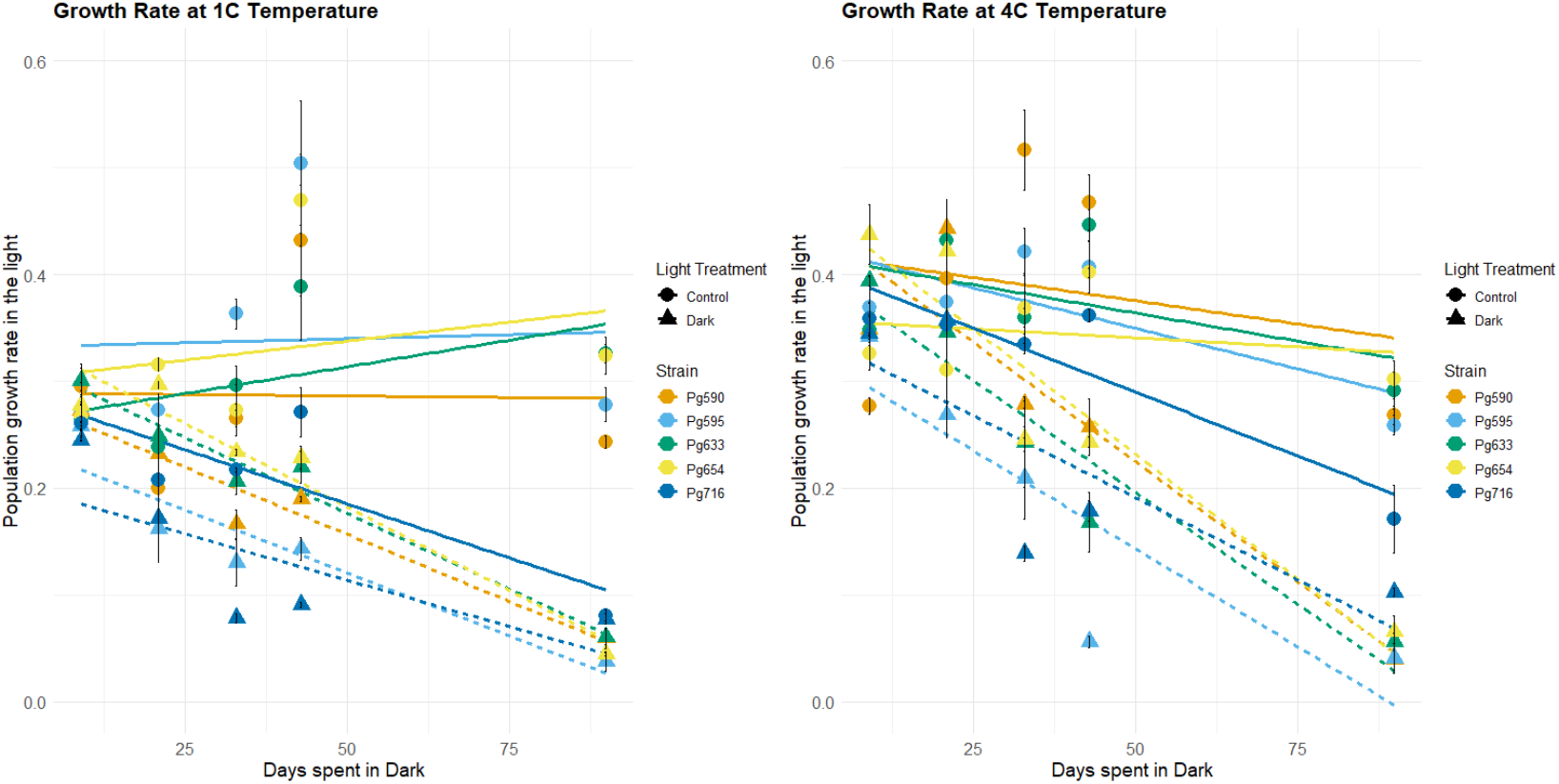
Growth rates upon re-illumination and re-inoculation of all temperature and light treatment combinations. a) shows the subsequent growth rates from control-light cultures (round points and solid lines) and dark cultures (triangles and dashed lines) in the 1°C treatment. b) shows the subsequent growth rates from control-light cultures and dark cultures in the 4°C treatment. Black bars at each point correspond to standard error. The colours of each line correspond to the 5 different strains of Porosira glacialis.

**Figure 4:**
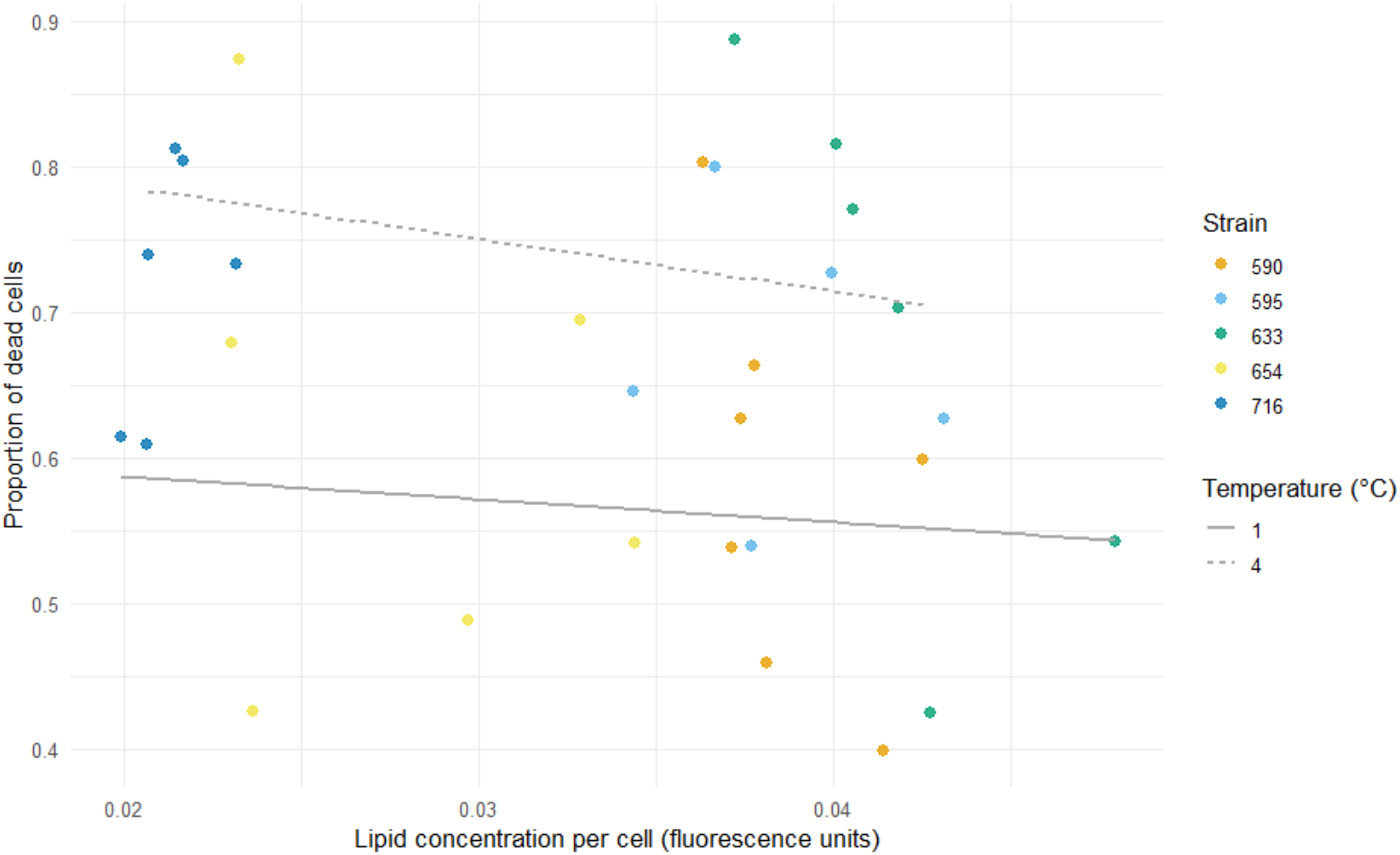
Mortality against Lipid concentration in Porosira glacialis cells. Solid line denotes cultures from the 1°C temperature treatment. Dotted line denotes cultures from the 4°C temperature treatment. All data concerns only cultures maintained in the dark.

**Figure 5.**
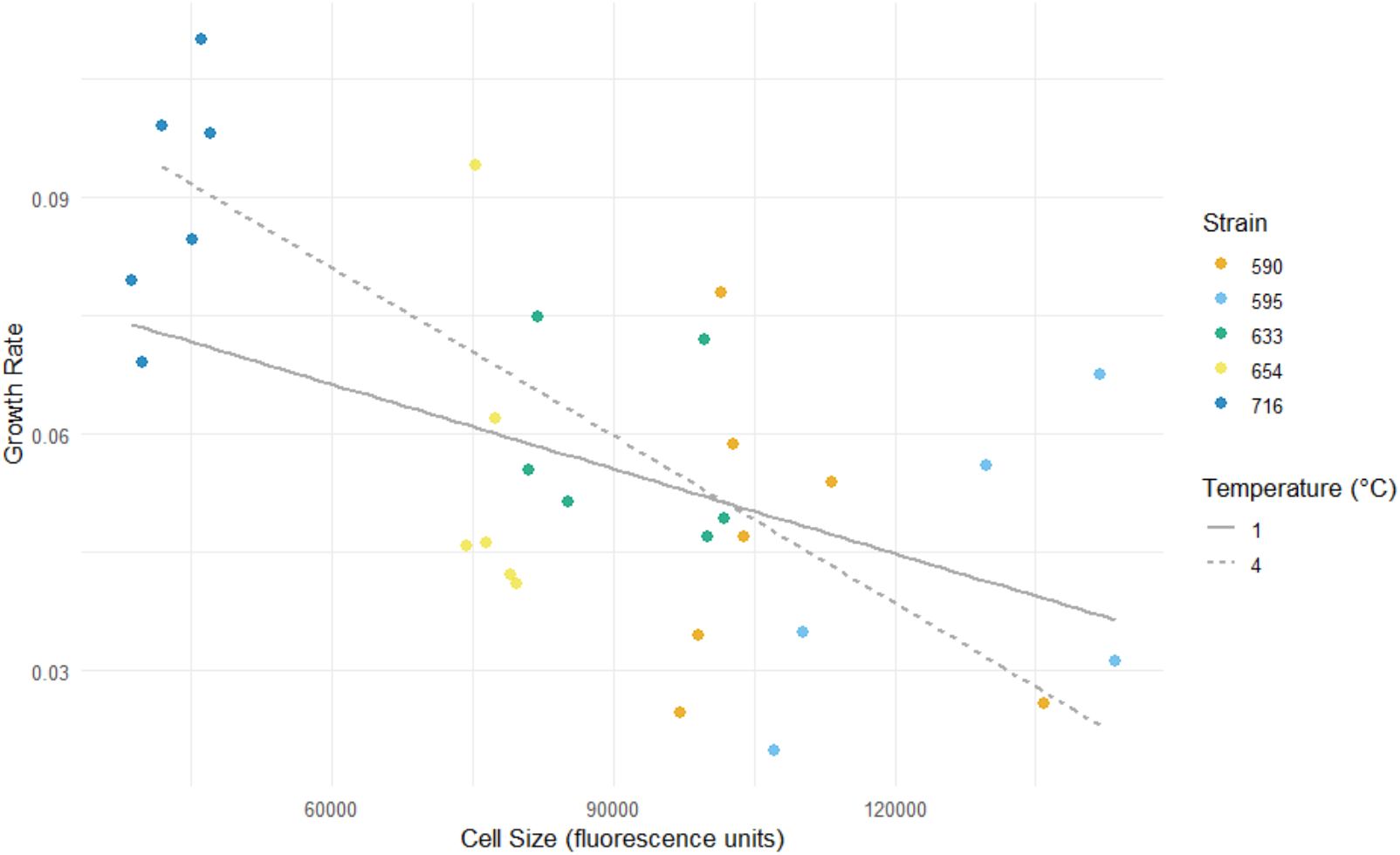
Growth rate as a function of median cell size (in fluorescence units) of Porosira glacialis strains. Solid line denotes cultures from the 1°C temperature treatment. Dotted line denotes cultures from the 4°C temperature treatment. All data concerns only cultures maintained in the dark.

Differences in cell survival and population growth rate following a period of darkness is exacerbated by the effect of temperature in this study. At the elevated temperature treatment (4°C) the average mortality was both higher and the growth rate on re-illumination was lower than in the low temperature treatment (1°C). Elevated temperature tends to increase metabolic activity (Bishop *et al*., 2022; McMinn and Martin, 2013), and it is plausible that higher mortality at 4°C is driven by cells being unable to lower metabolic activity as at warmer temperatures as much as they do at cooler ones, which in turn affects their dark survival (Joli *et al*., 2024; Zhang *et al*., 1998). The rank order of strains’ growth rates also changed between the 1°C and 4°C treatments, indicating that temperature not only exacerbates the effects of the polar night, but may independently change community composition since strains differ in their tolerance to warming (Bishop *et al*., 2022). This suggests that diatom community composition will not only be shaped by a strains’ ability to survive and resume growth following the polar night, but also at what temperatures these seasonal phenomena are experienced at. This is in line with scenario (c) in Figure 1.

Cell survival over prolonged dark periods might be mediated by their ability to enter a state of hypometabolism, and the consumption of cellular stores (Joli *et al*., 2024; Juchem *et al*., 2023; Schaub *et al*., 2017; Zhang *et al*., 1998). Based on this, we conducted a post-hoc experiment that measured lipid content after 3 months of darkness in the *P. glacialis* populations. Consistent with this, diatoms from the dark treatment in our experiment depleted their cellular lipid stores compared to diatoms in the light treatment (Schaub et al., 2017) (see S.I. Appendix 4, Figure S4). However, we found little evidence that lipid content explains the variation in strains’ ability to survive prolonged darkness or subsequent growth. Instead, temperature emerged as the best predictor for mortality in the lipid and mortality model, indicating that it is strain’s differential ability to tolerate elevated temperatures that drives variation in mortality in this experiment. Lipids correlated poorly with cell growth after re-illumination, and intraspecific differences in this effect is explained mainly by cell size (t = −3.454, p = 0.00618; 40% of variance in model), and temperature interaction with strains (explaining 16% of variance in the model). There was a weak correlation between lipid concentration and mortality (t = −1.986, p = 0.061682), this measurement was only made at a single timepoint and within a single species, so that variation in lipid content was low. This is a highly suggestive result given the low power of this experiment to detect it. Exploring the role of lipid content in mortality should be done using a greater range of lipid contents, and by studying both polar and nonpolar lipids.

Laboratory experiments are not, and do not draw their utility from, realism in terms of their representation of environments outside of the laboratory. Instead, the simplification of environments is what allows causal links to be made between specific environmental cues (in this case darkness) and organismal responses, which means that getting the environmental simplification right is vital. Recent evidence suggests that even minimal levels of light are sufficient to maintain baseline activity of photosystems (Hoppe, 2022) and that very faint light signals can be detected underwater, even during the polar night (Connan-McGinty *et al*., 2022). While using complete darkness to interrogate responses to polar nights has proven a useful starting point, comparing responses in complete darkness and under prolonged periods of extremely low light would further validate the increasing use of our and similar experimental setups.

To the best of our knowledge, this is one of very few studies that estimated mortality through direct counts of live and dead cells rather than proxy measurements such as cell concentrations or chlorophyll fluorescence (Reeves et al., 2011; van de Poll *et al*., 2020). To our knowledge, only Handy *et al*. (2024) directly assess cell integrity to investigate survival, and similarly find large variation in this between strains. This approach allows us to avoid potential confounding variables such as a final cell division in the dark (see S.I., Appendix 5, Figures S5, S6), varying rates of chlorophyll degradation between strains and treatments (Handy *et al*., 2024), or the preservation of intact but dead cells in cold systems. The overall trends based on this approach agree with wider research which supports decreasing survival over long periods of darkness (Handy *et al*., 2024; Joli *et al*., 2024; Morin *et al*., 2020; Reeves *et al*., 2011;), however, the magnitude of this was higher and more variable than previously reported. The magnitude, however, is not as interesting as the intraspecific variability we have found.

Our results demonstrate substantial intraspecific variation in both dark survival and population growth rate upon re-illumination in *Porosira glacialis*. This highlights the potential of the Polar night acting as a strong selective event between and within species of diatoms. The interaction between dark duration and temperature amplifies intraspecific variability in this study, indicating that warming has the potential to modify not only overall mortality and viability, but also the relative performance of strains and even community composition emerging from winter. Together, our findings emphasize the importance of considering intraspecific variability and demographic responses, rather than single “representative” strains, when predicting how Polar diatom populations and microalgal communities might respond to seasonal darkness and ongoing ocean warming.

## Acknowledgements

This project was supported by a British Phycological Society Research Grant to SC, and the University of Edinburgh MSc program in Ecology, Evolution and Biodiversity. We thank Shravan Raghu, Simin Gao and Katharin Balbirnie Cumming for laboratory assistance and training.

## Competing Interests

The authors declare no competing interests

### Author contributions

PM and SC designed the research. PM collected the data. PM and SC analysed the data. PM and SC wrote the manuscript.

## Data availability

Data and code can be found at https://github.com/PMrazek2001/Mrazek_PolarPhytoplankton

