## Supplemental Information for "Variation in survival and growth following prolonged darkness in a polar diatom species"

### Supporting Information

Appendix 1: Inoculation concentration of cultures into light and dark treatments

Table S1: Initial cell concentrations of culture replicates at inoculation into experimental treatments. Initial cell concentration was diluted by the Dilution factor (*culture:media) with the volume of culture and media specified in columns 7 & 8 for Dark treatments and 9 & 10 for Light treatment. Dilutions were calculated to achieve Final Cell Concentrations within the same order of magnitude and as similar as possible, however strain 595 had a consistently lower cell concentration and growth rate. Note also that a further 1 in 10 dilution is applied to cultures going into the light treatment to allow for growth.

| Sample ID | Temperature | Strain | InitialCellConc (cells/ml) | Dilution factor* | FinalCellConc | VolCulture Dark | VolMedia Dark | VolCulture Light | VolMediaLight | FinalVol |
|---|---|---|---|---|---|---|---|---|---|---|
| Pg654L | 1 | 654 | 6388 | 2/3 | 4258.2408 | 33.33333333 | 16.66666667 | 3.333333333 | 46.66666667 | 50 |
| Pg716L | 1 | 716 | 11013.8 | 2/5 | 4405.52 | 20 | 30 | 2 | 48 | 50 |
| Pg595L | 1 | 595 | 2620.4 | Na | 2620.4 | 50 | 0 | 5 | 45 | 50 |
| Pg590L | 1 | 590 | 4566.6 | Na | 4566.6 | 50 | 0 | 5 | 45 | 50 |
| Pg633L | 1 | 633 | 4344.4 | Na | 4344.4 | 50 | 0 | 5 | 45 | 50 |
| Pg654H | 4 | 654 | 6805.5 | 3/5 | 4083.3 | 30 | 20 | 3 | 47 | 50 |
| Pg716H | 4 | 716 | 15958.3 | 1/4 | 3989.575 | 12.5 | 37.5 | 1.25 | 48.75 | 50 |
| Pg595H | 4 | 595 | 2212.9 | Na | 2212.9 | 50 | 0 | 5 | 45 | 50 |
| Pg590H | 4 | 590 | 3731.5 | Na | 3731.5 | 50 | 0 | 5 | 45 | 50 |
| Pg633H | 4 | 633 | 6866.6 | 3/5 | 4119.96 | 30 | 20 | 3 | 47 | 50 |

Appendix 2: Experimental Design

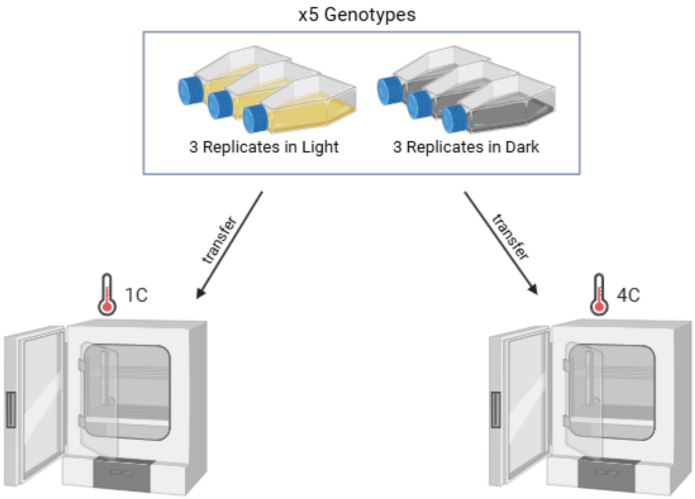


*Figure S1: Experimental design. Three replicates of each of 5 genotypes were either wrapped in 3 layers of tinfoil (to create a dark environment) or left in the light (at 25-35 micromol/m/s). Then 3 replicates of each light treatment were placed in either a 1C incubator or a 4C incubator. Figure created using BioRender.com*

Figure S2:

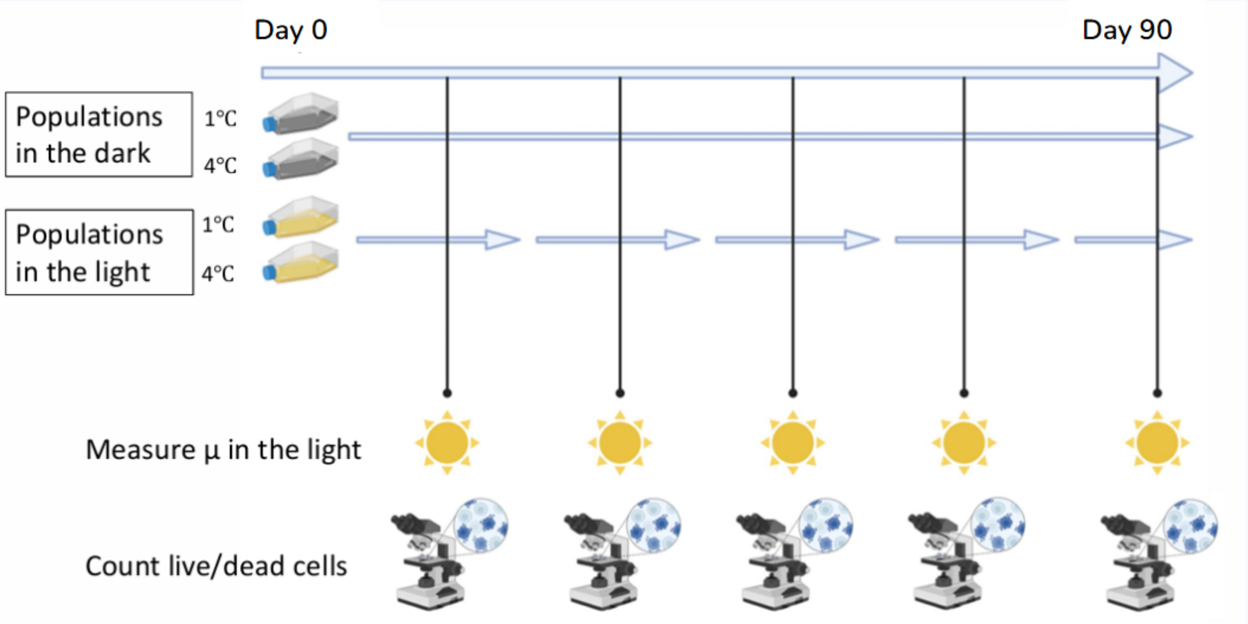


*Figure S2: Sampling timeline design. Cultures were sampled for growth rate on re-illumination and mortality at 5 timepoints. Growth rate re-illumination timepoints occurred on days 9, 21, 33, 43 and 90 of the experiment, with day 0 being when cultures were placed in their respective experimental treatments. Mortality was sampled on days 12, 22, 36, 50 and 85 of the experiment. Sampling days for these two assays were staggered to ensure feasibility.* Figure created using BioRender.com

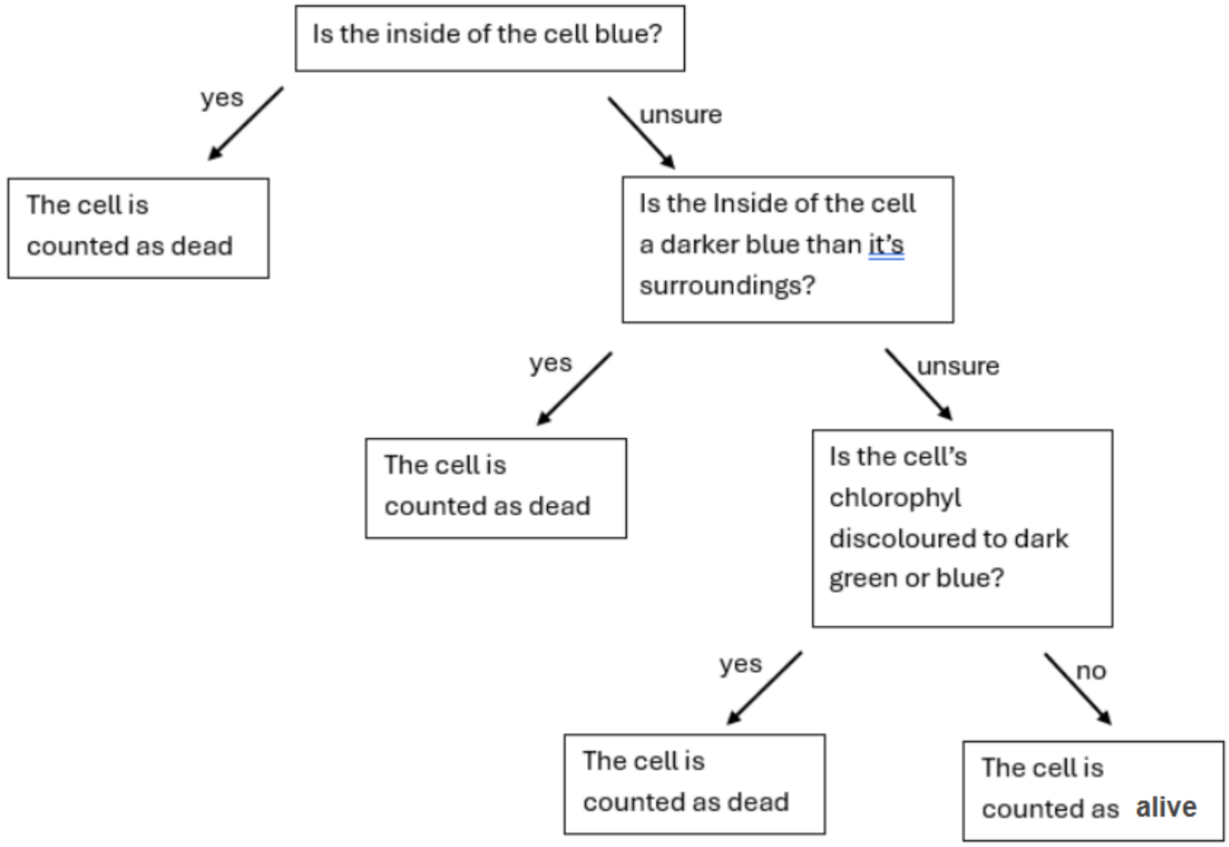


*Figure S3: Flow chart of decision process to determine whether a cell is live or dead.*

Appendix 3: FlowJo Gating Information and Statistical Models:

Flow cytometry data was gated in FlowJo following certain criteria to exclude organic matter that was not diatom cells. We gated the data to exclude chlorophyll-negative events, ensuring we deal with live cell data rather than debris. We also gated the data by size, excluding events at extreme small or large ends of the forward scatter (fst - which is a proxy for spherical cell volume) spectrum.

We tested four lipid hypotheses for both mortality and growth rate:

H0 - lipid content does not explain variation in mortality/growth rate upon re-illumination

H1 - initial lipid content best explains variation in mortality/growth rate

H2 - Final lipid content best explains variation in mortality/growth rate

H3 - Lipid loss best explains variation in mortality/growth rate.

The fit of each model was evaluated using AIC and R2 and residuals meeting model assumptions.

Mortality Hypothesis models:

H0:

Prop. dead = Intercept + β1*median size + β2*strain + β3*temperature + β4*median size*strain + β5*median size*temperature + β6*strain*temperature + Error

R2 = 61.85%

AIC = -53.61882

H1:

Prop. dead = Intercept + β1*initial lipid content + β2*strain + β3*temperature + Error

R2 = 46.54%

AIC = -44.50633

H2:

Porp.dead = Intercept + β1*lipid concentration + β2*strain + β3*lipid concentration*strain + β4*temperature

R2 = 54.27%

AIC = -46.92257

H3:

*Prop.dead = Intercept + β1*lipid fold change + β2*Strain + β*Temperature + Error*

R2 = 48.24%

AIC = -45.4789

Note that the presented models are only ones that showed best fit for each hypothesis, determined based on R2, AIC and residual plots.

We proceeded with model H2, which fits the proportion of dead cells as a function of the lipid concentration per median cell spherical volume (in fluorescence units), strain and temperature. This model had the best combination of R2, AIC and residuals plots of the considered models.

Growth rate Hypothesis Models:

H0: GH0a <- lm(growthrateB ~ Size_Median*Strain*Temp, data = BDDark)

*Growth rate = β1*median size + β2*Strain + β3*temperature + β4*median size*Strain + β5*median size*temperature + β6*Strain*temperature + β7*median size*Strain*temperature + Error*

R2 = 79.28%

AIC = -177.6802

H1: GH1b <- lm(growthrateB ~ mean_lightBD*Temp + Strain, data = BDDark)

*Growth rate = Intercept + β1*initial lipid content + β2*temperature + β3*initial lipid content*temperature + β4*Strain*

R2 = 45.46%

AIC = -148.9965

H2: GH2 <- lm(growthrateB ~ Bodipy_Median*Strain*Temp, data = BDDark)

*Growth rate = Intercept + β1*median lipid content + β2*Strain + β3*temperature + β4*median lipid content*Strain + β5*median lipid content*temperature + β6*strain*temperature + β7*median lipid content*Strain*temperature + Error*

R2 = 59.11%

AIC = -157.2883

H3: GH5b <- lm(growthrateB ~ fold_change*Temp*Size_Median + Strain, data = BDDark)

*Growth rate = Intercept +* $\beta$*1*lipid fold change +* $\beta$*2*temperature +* $\beta$*3*median size +* $\beta$*4*lipid fold change*temperature +* $\beta$*5*lipid fold change*median size +* $\beta$*6*temperature*median size +* $\beta$*7*lipid fold change*temperature*median size +* $\beta$*8*Strain + Error*

$R2 = 63.03\%$

AIC = -158.6839

Note again, that the presented models are only ones that showed best fit for each hypothesis, determined based on R2, AIC and residual plots.

We proceeded with model H0, as this model had the highest R2 while the AIC was not much worse than other models and residuals plots were in line with linear model assumptions. An argument can be made for model H3, however due to the limited power in this study and potential confounding effects between lipid fold change and cell size, we opted for model H0, further considering that cell size was a consistently significant explanatory variable for growth rate upon re-illumination.

Appendix 4: Lipid Results with both light and dark treatments

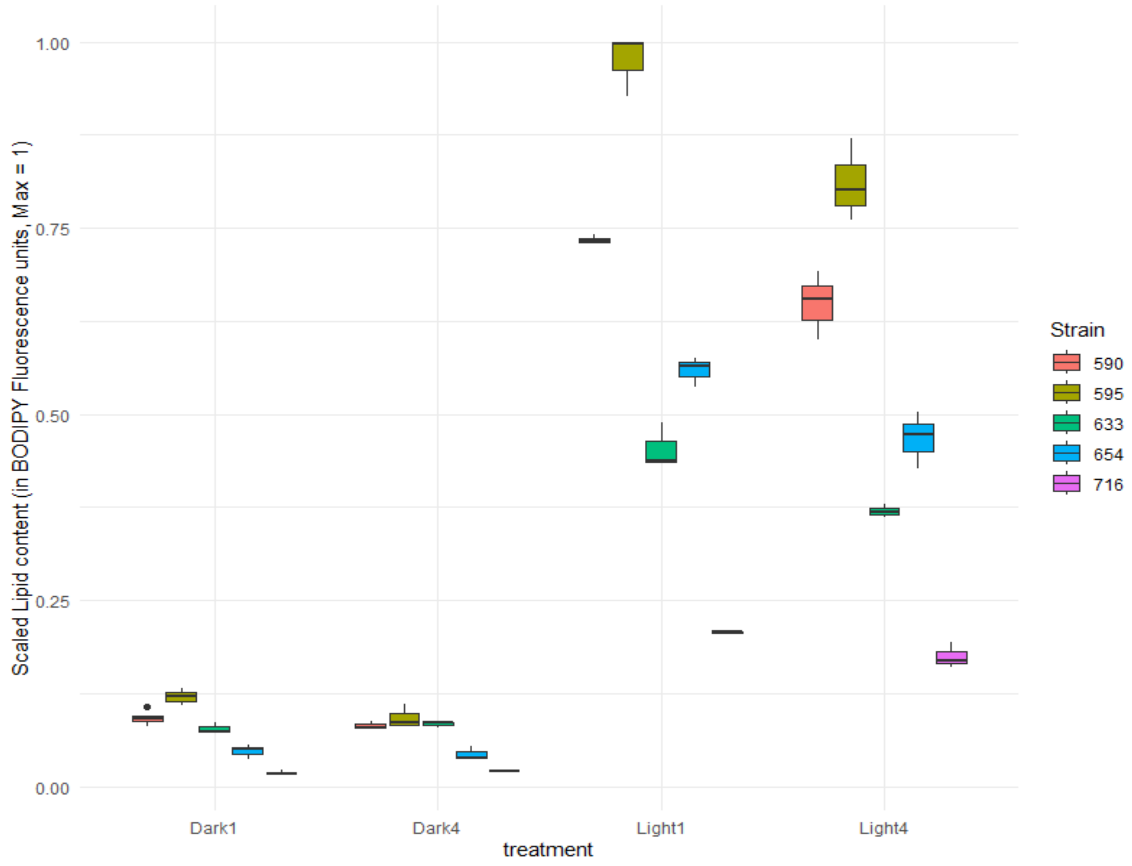


*Figure 4: Lipid content, in scaled Bodipy fluorescence units, in the 4 experimental treatments. From left to right: Dark treatment 1C temperature; Dark treatment 4C temperature; Light treatment 1C temperature; Light treatment 4C temperature. There is a clear disparity between Light and Dark treatments with further hierarchy between strains within each treatment. Different colors indicate different strains. Horizontal line within boxes shows the median and the whiskers are 75$^{th}$ percentile ranges.*

Analysing lipid content across both light and dark treatments, using the model below, shows a significant difference in polar lipid content between the two Light treatments ($F = 440.3932$, $p \ll 0.001$, $DF = 56$), and temperature treatments ($F = 9.9415$, $p = 0.002597$, $DF = 56$), and includes an interaction between temperature and light ($F = 7.9824$, $p = 0.006534$, $DF = 56$), with temperature exacerbating lipid loss in both light treatments, but being more severe in the dark (Figure S4). This is in line with our expectations and wider research. Further lipid analysis was limited to data from the dark treatments to avoid confounding effects from lipid content and mortality in the light treatments.

*Lipid concentration = Intercept + β1*Light treatment + β2*temperature + β3*Light treatment*temperature + Error*

Appendix 5: Cell size and concentration

Our cell concentration data sampled throughout the experiment and cell size (here represented by a proxy of median cell spherical volume, fsc, from flow-cytometry measurements) indicate a possible cell division in the dark. Notably for strain Pg633, cell concentration increases between timepoint 1 and 2 (Figure S6). Though some sampling error might increase variability, coupled with a decreased cell size in the dark vs in the light (Figure S5), this at least suggests a final cell division occurring between 2 and 4 weeks after beginning the dark treatment is possible for some of the strains used here.

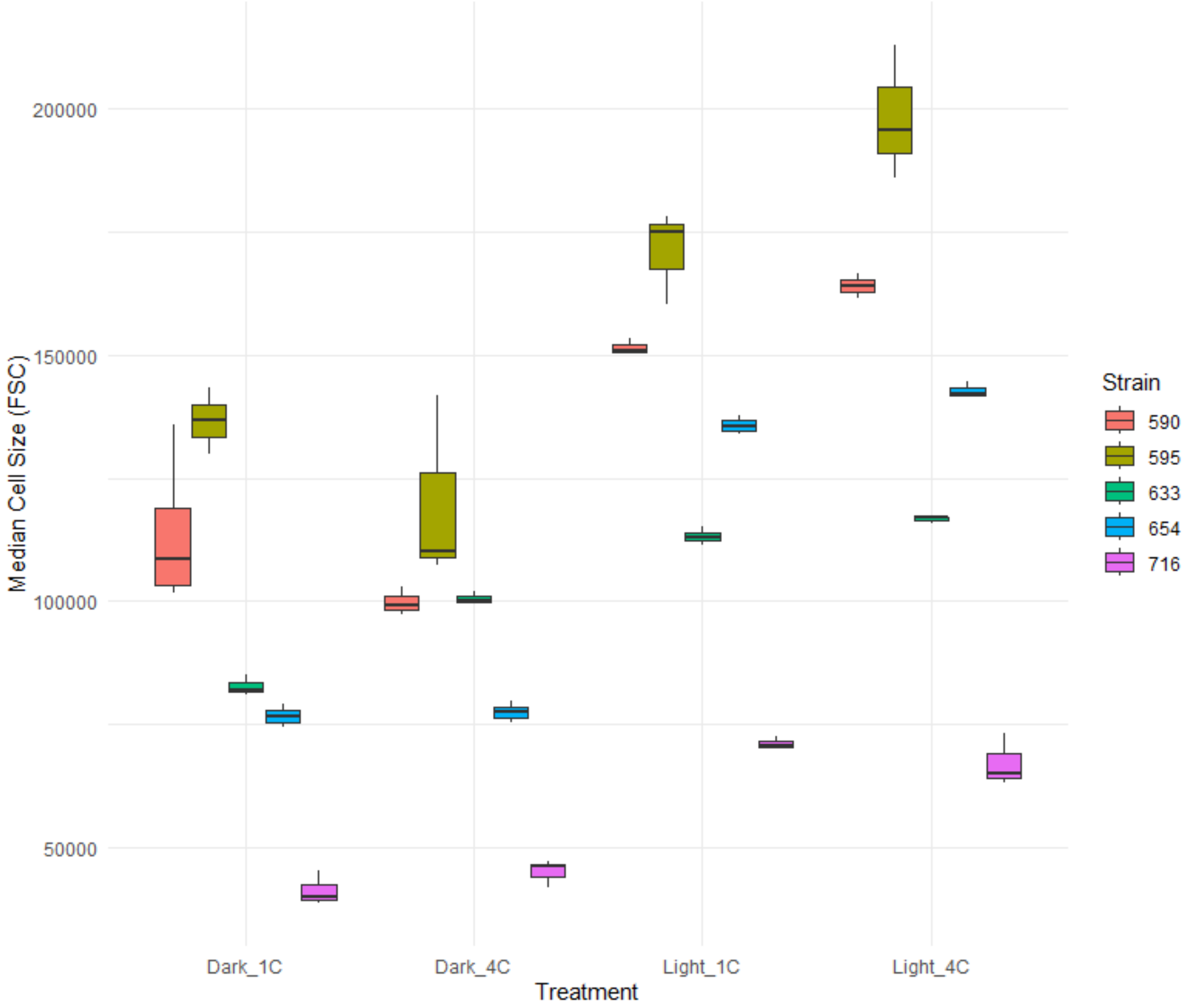


*Figure S5: Median cell size (as median cell spherical volume in FSC units) of* Porosira glacialis *following 3 months of experimental treatment (x), separated by strain. Different colors indicate different strains. Horizontal line within boxes shows the median and the whiskers are 75<sup>th</sup> percentile ranges. Median cell size appears to be larger in the light treatment vs the dark treatment.*

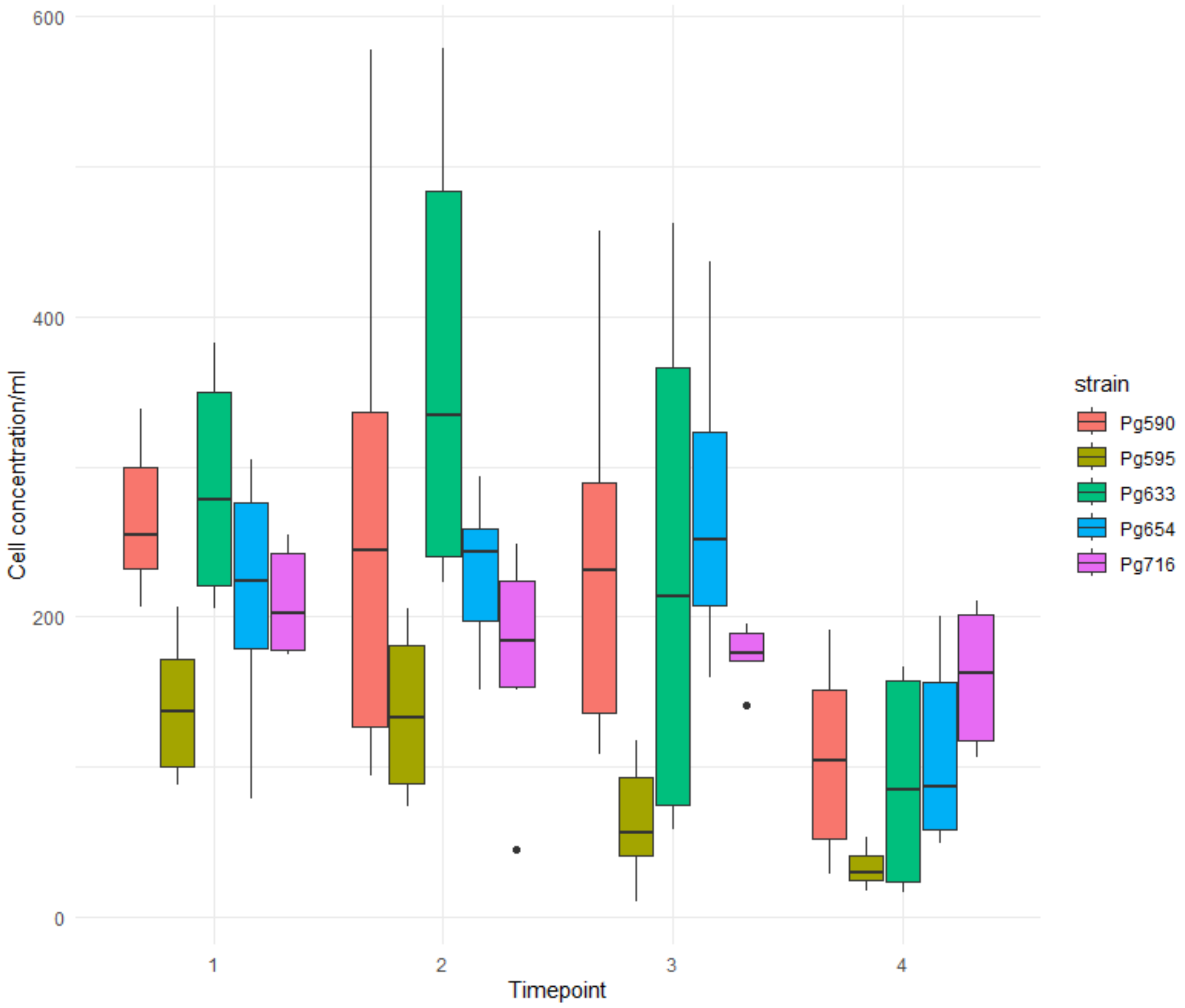


*Figure S6: Cell concentration at each timepoint, separated by strain. Different colors indicate different strains. Horizontal line within boxes shows the median and the whiskers are 75th percentile ranges.*
